# HTLV-1 infected T cells cause bone loss via small extracellular vesicles

**DOI:** 10.1101/2024.02.29.582779

**Authors:** Nitin Kumar Pokhrel, Amanda Panfil, Haniya Habib, Sham Seeniraj, Ancy Joseph, Daniel Rauch, Linda Cox, Robert Sprung, Petra Erdmann Gilmore, Qiang Zhang, R Reid Townsend, Lianbo Yu, Ayse Selen Yilmaz, Rajeev Aurora, William Park, Lee Ratner, Katherine N Weilbaecher, Deborah J Veis

## Abstract

Adult T cell leukemia (ATL), caused by infection with human T cell leukemia virus type 1 (HTLV-1), is often complicated by hypercalcemia and osteolytic lesions. Therefore, we studied the communication between patient-derived ATL cells (ATL-PDX) and HTLV-1 immortalized CD4+ T cell lines (HTLV/T) with osteoclasts and their effects on bone mass in mice. Intratibial inoculation of some HTLV/T lead to a profound local decrease in bone mass similar to marrow-replacing ATL-PDX, despite the fact that few HTLV/T cells persisted in the bone. To study the direct effect of HTLV/T and ATL-PDX on osteoclasts, supernatants were added to murine and human osteoclast precursors. ATL-PDX supernatants from hypercalcemic patients promoted formation of mature osteoclasts, while those from HTLV/T were variably stimulatory, but had largely consistent effects between human and murine cultures. Interestingly, this osteoclastic activity did not correlate with expression of osteoclastogenic cytokine RANKL, suggesting an alternative mechanism. HTLV/T and ATL-PDX produce small extracellular vesicles (sEV), known to facilitate HTLV-1 infection. We hypothesized that these sEV also mediate bone loss by targeting osteoclasts. We isolated sEV from both HTLV/T and ATL-PDX, and found they carried most of the activity found in supernatants. In contrast, sEV from uninfected activated T cells had little effect. Analysis of sEV (both active and inactive) by mass spectrometry and electron microscopy confirmed absence of RANKL and intact virus. Viral proteins Tax and Env were only present in sEV from the active, osteoclast-stimulatory group, along with increased representation of proteins involved in osteoclastogenesis and bone resorption. sEV injected over mouse calvaria in the presence of low dose RANKL caused more osteolysis than RANKL alone. Thus, HTLV-1 infection of T cells can cause release of sEV with strong osteolytic potential, providing a mechanism beyond RANKL production that modifies the bone microenvironment, even in the absence of overt leukemia.

## 1 Introduction

Adult T cell leukemia (ATL), an aggressive, treatment-refractory malignancy, occurs in 2-5% of patients infected with the retrovirus HTLV-1, often decades after initial infection^1^. Clinical presentation often includes hypercalcemia, a life-threatening complication, due to release of the mineral from bone^2,3^. Previous studies of ATL have shown expression of soluble factors such as RANKL, MIP1α, and TNF ^3,4,5,6,7,8,9^ which directly enhance formation of osteoclasts, the myeloid-lineage polykaryons responsible for both physiologic and pathologic bone resorption. ATL cells also can produce PTHrP, a growth factor associated with many forms of hypercalcemia of malignancy, that exerts its effects indirectly, via the osteolineage^4,5,10^. Bone is also a rich source of matrix-stored growth factors such as transforming growth factor β (TGF-β) and insulin like growth factor 1 (IGF1) ^11,12^ which can interact with tumor cells in bone, affecting tumor growth and dormancy. In addition, recent studies have shown that tumor cells in non-skeletal primary sites communicate with the bone microenvironment, preparing it to be receptive to local tumor cell dissemination and growth, i.e. formation of a premetastatic niche^13,14^. It has been shown, using mouse models, that alterations to the bone microenvironment are also important for hematologic malignancies^15,16^. However, factors favoring transition to malignancy following chronic viral infection, such as with HTLV-1, are largely unknown. Here we investigate communication from HTLV-1-infected T cells to osteoclasts, which may represent a first step.

A number of model systems have been used to study the osteolysis associated with ATL, employing tumor cell lines, HTLV-1 infection of humanized mice, and transgenic mice^17^. We have previously published that established ATL cell lines have multiple phenotypes (osteolytic, osteoblastic and mixed) when implanted locally into bone, and their expression of soluble osteoclastogenic factors is not uniform^4^. HTLV-1 infection of humanized mice or implantation of patient-derived ATL cells (ATL-PDX) in mice cause systemic bone loss, but these models have rapidly progressive leukemia or polyclonal lymphoproliferation associated with systemic inflammation, confounding the analysis of bone pathophysiology and evaluation of early events in the transition to malignancy^6^. Furthermore, human anti-RANKL antibody provides incomplete protection from bone loss in the infection model^6^. We have also studied the bone phenotype of transgenic mice expressing the HTLV-1 oncogenes Tax and Hbz under the granzyme B (GzmB) promoter^18,19^. Tax, a coactivator of CREB binding proteins, activates many viral and cellular genes and is capable of inducing inflammation as well as transformation of multiple cell types. Tax^GzmB^ transgenic mice develop subcutaneous tumors composed of cells that are fully transformed and transplantable but have characteristics of NK cells rather than T cells. Hbz, a helix-basic loop zipper protein, is expressed at late stages in ATL and regulates proliferation as well as expression of inflammatory factors including RANKL^6^. Hbz^GzmB^ transgenic mice develop lymphoproliferation of both CD4 and CD8 positive cells, although ATL cells are almost always CD4 positive. In both transgenic lines, mice develop osteolysis and hypercalcemia, with a long latency. Thus, although useful, each of these mouse models has key differences in pathophysiology compared to human ATL.

In addition to releasing the many soluble factors previously studied in the context of osteolysis, HTLV-1 infected cells, like uninfected cells, release small extracellular vesicles (sEV), membrane-bound vesicles enriched in multiple biological cargos including proteins, metabolites, miRNAs and mRNAs^20^. Initially considered mere waste-carrying vehicles, sEV are now recognized as important mediators of intercellular communication in homeostasis and pathologic conditions including cancer^21^. An early study of sEV released by HTLV-1 infected cells demonstrated transfer of the oncogene Tax^22^. These sEV increase viral spread and disease progression during HTLV-1 infection by promoting cell to cell contact^23,24^. More generally, sEV from tumor cells are believed to provide signaling cues for initiation of metastasis, immune evasion and preparation of the pre-metastatic niche^20,21^. Studies with cell lines have demonstrated direct osteoclastogenic effects of breast and prostate tumor-derived sEV^25,26^, and indirect effects via enhanced RANKL production by osteolineage cells in multiple myeloma^27^. sEV from ATL cell lines and PDX were shown to modulate bone marrow stromal cell proliferation and gene expression, suggesting that they could modify the bone microenvironment^28^. However, to date there are no published studies examining effects of sEV on bone *in vivo*, in the context of ATL or HTLV-1 infection.

In this study, we utilized a panel of HTLV-1 infected T cell lines to study their interactions with osteoclasts in culture as well as in mouse models, in comparison with ATL-PDX. We found that sEV from both HTLV-1-infected T cell lines and ATL-PDX can promote *in vitro* osteoclastogenesis and *in vivo* bone resorption. In addition, these cell lines lead to consistent *in vivo* osteolysis in mice independent of RANKL/OPG expression.

## 2 Methods

### 2.1 Animals and cells

6-week-old NOD-*Prkdc*^*em26Cd52*^*Il2rg*^*em26Cd22*^/NjuCrl (NCG) mice (Charles River, Wilmington, MA) were used for *in vivo* experiments with cell and sEV implantation. The numbers of male and female mice were matched in all the experimental groups. Mice were kept with *ad libitum* access to fresh chow and water in a pathogen-free temperature-controlled barrier facility. Mice were monitored daily by staff of the Division of Comparative Medicine. 6-8 week old immune competent C57Bl/6 female mice were used for bone marrow macrophage (BMM) isolation. Human peripheral blood mononuclear cells (hPBMCs) were isolated by density gradient centrifugation of the blood from anonymous healthy donors. All the animal experimental protocols were approved by Washington University School of Medicine’s Animal Studies Committee (approvals 19-1059 and 22-0339).

### 2.2 Generation of HTLV/T cells

HTLV/T were created by co-culturing lethally irradiated HTLV-1 producer cells with freshly isolated hPBMCs. After a period of 10-14 weeks, HTLV-1-immortalized CD4+ T-cells were established and were maintained in RPMI with 20% FBS, 1% pen/strep and 20 U/mL human IL-2. After establishment of cell lines, they were validated for surface expression of CD3 and CD4 by flow cytometry^29,30^.

### 2.3 BMM isolation and osteoclast differentiation

Bone marrow was harvested from dissected femurs and tibiae of 2 month old C57Bl/6 mice via centrifugation at 12000 rpm for 2 minutes. Resuspended marrow was passed through 40µM filter and cells cultured in alpha MEM (Sigma, M0894) containing 10% FBS (Gibco), 100 IU/mL penicillin/streptomycin, and 1:10 dilution of CMG 14-12 cell supernatant containing an equivalent of 100 ng/ml Macrophage-Colony Stimulating Factor (M-CSF) for 5-7 days. The resulting expanded BMMs were plated 10,000 cells/well in 96-well plates. Purified GST-RANKL^31^ at 10 ng/ml was added to alpha-MEM containing 10% FBS, 100 IU/ml penicillin/streptomycin, and a 1:50 dilution of CMG14-12 cell supernatant^32^. The medium was changed every alternate day. TRAP staining was performed after fixation with 4% paraformaldehyde (Polysciences, Warrington, PA) and 0.1% Triton X-100 in PBS according to the manufacturer’s instructions (Sigma, 387A). The plates were left to air dry and imaged using a light microscope (Olympus BX41, Tokyo, Japan).

### 2.4 Resorption Assay

Expanded BMMs (5×10^5^ cells) were seeded on 6 well plates and treated with 50 ng/mL of RANKL for 48h for the generation of pre-osteoclasts. These were lifted and plated on bovine bone slices with or without 10% supernatant together with 10 ng/mL of RANKL and 1:50 dilution of CMG14-12 cell supernatant. After 5 days, bone slices were incubated in 0.5 NaOH for 30s, then rinsed in PBS. Cells were removed using a cotton swab and bone slices were incubated with 20 μg/ml peroxidase-conjugated wheat germ agglutinin in PBS (Sigma, 61767) for 30 min. Bone slices were washed three times in PBS and incubated with DAB Chromogen kit (#BDB20004H, Biocare Medical, Biocare Medical, Pacheco, CA) at 37°C for 30 min. Air dried slices were imaged by light microscope as above.

### 2.5 In vivo micro-computed tomography (microCT)

The right tibia of mice was scanned by microCT *in vivo* (VivaCT 40, Scanco, Brüttisellen, Switzerland) at 10.5 μm resolution (70 kVp, 114 mA, 8 W, 100 ms integration time) at 2 or 4 week intervals. Cancellous bone parameters were measured in a 1 mm region distal to the end of the tibial growth plate.

### 2.6 Ex vivo micro-CT

The calvaria of mice was scanned by microCT ex vivo (MicroCT 50, Scanco) at 7 μm resolution (70 kVp, 57uA, 4 W, 700 ms integration time). Calvarial BV/TV encompassed 200 slices at each region of interest.

### 2.7 Genomic DNA quantitative PCR

Genomic DNA was prepared from flushed marrow of cell line-injected tibiae using DNeasy Blood & Tissue Kit (#69504, Qiagen, Hilden, Germany,). A non-injected mouse bone marrow was used as a negative control. A standard curve was created by spiking a known number of human cells in increasing percentage into a fixed number of mouse bone marrow cells. Quantitative PCR program was run on ABI QuantStudio 3 with iTaq Universal SYBR Green Supermix (#1725121, Bio-Rad, Hercules, CA,) using human *Il6* and mouse *B2m*, using species-specific regions (see Table 1). Each reaction proceeded at 50 °C for 2 min, 95 °C for 10 min, and then 40 cycles of 95 °C for 15 s and 60 °C for 1 min. Relative amount of human DNA was calculated as 2^-^(hIL6 CT -mB2M CT)^. Percentage of human cells was calculated by plotting values against the standard curve.

### 2.8 sEV Isolation

1 million HTLV/T cells were cultured in RPMI supplemented with 20% exosome-depleted FBS, 1% Penicillin/Streptomycin and 20 IU human IL-2 (Roche Diagnostics, Mannheim, Germany) for 5d. FBS was depleted of exosomes by centrifugation of heat inactivated serum at 2000 rpm for 15 min at room temperature, followed by another centrifugation of supernatant at 17.5K RPM (35000Xg) for 15 min at 4C. Vesicles were isolated using either of two commercial exosome isolation reagents (Total Exosome Isolation Reagent #4478359, Thermo Fisher Scientific; ExoQuick Exosome Isolation #EXOTC-10A, SBI, Palo Alto, CA) using the manufacturers’ protocols. The precipitated sEV were resuspended in PBS and stored at -80°C. No significant differences were observed in the activity of sEV between these kits. Isolated sEV were analyzed for size and distribution using NanoSight NS300 (Malvern Panalytical, Worcestershire, UK).

### 2.9 sEV uptake Assay

BMMs were grown on coverslips. sEV were labelled with PKH 26 (MINI 26, Sigma, St. Louis, MO) following the manufacturer’s protocol. The cells were treated with labelled sEV for 4 h. Images were captured using Leica DMi8 automated inverted microscope equipped with ACS APO 20x/0.60 Lens (Leica Microsystems, Wetzler, Germany).

### 2.10 RNA isolation and reverse transcription quantitative real-time PCR

After cell lysis with TRIzol (Life Technologies), solution was subjected to phenol:chloroform extraction followed by centrifugation at 12,000 g, and the aqueous layer was harvested. An equal volume of 70% ethanol was added, and the remaining RNA isolation was done using the NucleoSpin RNA II Kit (Clontech Laboratories, Palo Alto, CA; 740955.50). 1 μg RNA was transferred into the cDNA Ecodry Premix Kit prior to the quantitative PCR program being run on ABI QuantStudio 3 with iTaq Universal SYBR Green Supermix (Bio-Rad, Hercules, CA, 1725121). Each reaction proceeded at 50 °C for 2 min, 95 °C for 10 min, and then 40 cycles of 95 °C for 15 s and 60 °C for 1 min. Relative expression was calculated as 2^-^(target CT -control CT)^. The primer sequences used are listed in Supplementary Table 1.

### 2.11 Intratibial injection of cells

Mice were kept under anesthesia with isoflurane and the right knee joint was exposed and sterilized. A 27G needle was inserted through knee joint to make a hole, 2.5×10^6^ cells in 10 *μ*l PBS were injected and the skin incision was sutured. Buprenorphine ER was given as a pain reliever.

### 2.12 Calvarial Injection of sEV

NCG mice (6 weeks of age) were anesthetized with isoflurane and injected subcutaneously over the calvarium once per day. For the first 2 days, all mice received 50 μg RANKL, and next 3 days 25 μg RANKL along with either PBS or sEV from HTLV/T C8. Calvaria were harvested for microCT 3 days after the last treatment.

### 2.13 Transmission electron microscopy

sEV were fixed with 1% glutaraldehyde (Ted Pella Inc., Redding CA) and allowed to absorb onto freshly glow discharged formvar/carbon-coated copper grids (200 mesh, Ted Pella Inc.) for 10 min. Grids were then washed two times in dH2O and stained with 1% aqueous uranyl acetate (Ted Pella Inc) for 1 min. Excess liquid was gently wicked off and grids were allowed to air dry. Samples were viewed on a JEOL 1200EX transmission electron microscope (JEOL USA, Peabody, MA) equipped with an AMT 8-megapixel digital camera (Advanced Microscopy Techniques, Woburn, MA).

### 2.14 Small RNA Sequencing

RNA was isolated from precipitated sEV using Sera Mir (Exosome RNA Column Purification Kit #RA808A-1, SBI). Total RNA integrity was determined using Agilent Bioanalyzer or 4200 Tapestation. Library preparation was performed with 1ug of total RNA with a Bioanalyzer RIN score greater than 8.0. Full library preparation was performed using the TruSeq Small RNA kit (Illumina, San Diego, CA) per manufacturer’s protocol. Fragments were sequenced on an Illumina NextSeq-550 using single reads extending 75 bases. Basecalls and demultiplexing were performed with Illumina’s bcl2fastq software with a maximum of one mismatch in the indexing read. RNA-seq reads were then aligned to the Ensembl release 101 primary assembly with STAR version 2.7.9a1. Gene counts were derived from the number of uniquely aligned unambiguous reads by Subread: feature Count version 2.0.32. Isoform expression of known Ensembl transcripts were quantified with Salmon version 1.5.23. Sequencing performance was assessed for the total number of aligned reads, total number of uniquely aligned reads, and features detected. The ribosomal fraction, known junction saturation, and read distribution over known gene models were quantified with RSeQC version 4.04. Despite using 8-10-fold more supernatant as starting material for sEV preparation, we were unable to obtain enough RNA from sEV without effect on osteoclasts.

### 2.15 Proteomics

sEV isolated from human HTLV/T cell cultures with Total Exosome Isolation Reagent (#4478359, Thermo Fisher Scientific) were solubilized in SDS buffer, digested, and quantified, then analyzed using trapped ion mobility time-of-flight mass spectrometry. Comprehensive methodology and data processing is provided in the supplemental text.

### 2.16 Histology

Mouse calvaria were fixed in 10% neutral buffered formalin for 24 h followed by decalcification in 14% Free Acid EDTA (pH 7.2) for 5d. 7μM sections were prepared and subjected to TRAP staining. Number of osteoclasts per bone perimeter (N.Oc/B.Pm) was calculated using Image J.

### 2.17 Statistical Analysis

All statistics were computed using SAS software (Version 9.4, SAS Institute Inc., Gary, NC, USA), GraphPad Prism software (Version 10.1.1 (270), GraphPad Software, Inc., La Jolla, CA, USA). Values of p < 0.05 were considered significant and data are presented as mean ± SD. For two group comparisons, a student’s, paired or unpaired, two-tailed t-test was used. For multiple group comparisons, a 1-way ANOVA followed by Tukey’s multiple comparisons test was performed. For repeated measure group comparisons, linear mixed-effects models were used followed by Tukey’s method for multiplicity adjustment. Specific statistical tests and sample sizes are indicated in the respective figure legends.

## 3 Results

### 3.1 HTLV-1 infected T cells cause bone loss in mice even in the absence of overt leukemia

Previously, we reported that established ATL cell lines, directly inoculated into bone, have variable ability to cause *in vivo* osteolysis^4^. In contrast, intraperitoneal administration of patient derived cell lines (ATL-PDX) or infection of humanized mice with HTLV-1 leads to massive systemic bone loss, but these mice also have rapidly progressive leukemia or lymphoproliferation^6^ making it difficult to discern whether the bone loss is directly caused by HTLV-1 infection or is a consequence of high systemic T cell burden. To address this, we locally injected RB, an ATL-PDX derived from a hypercalcemic patient, in one tibia of non-humanized NCG mice and monitored bone density by vivaCT. As expected, RB caused massive local bone loss by four weeks (Figure 1a), but these mice also developed lethal systemic leukemia with concomitant splenomegaly and complete bone marrow replacement within 6 weeks (Fig 1b, RB cell line).

**Figure 1.**
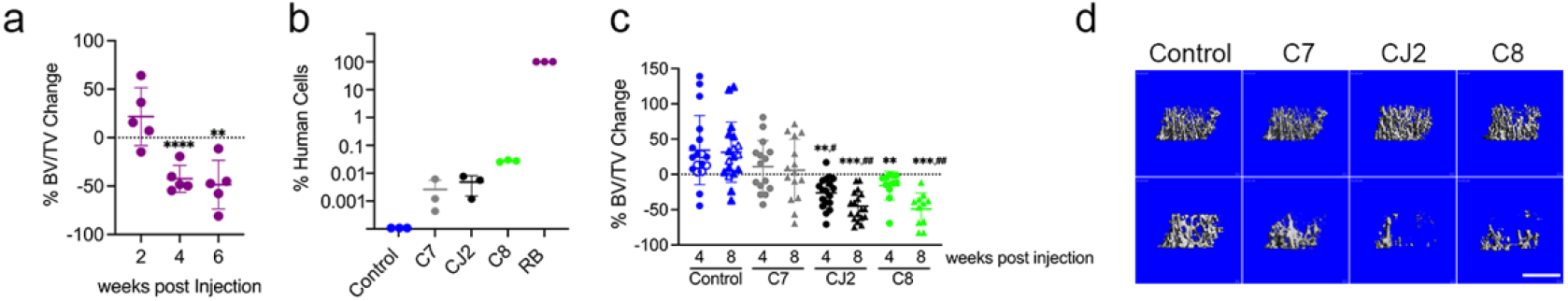
Implantation of HTLV-1 infected T cell lines into bone causes osteolysis even in the absence of overt leukemia. a) NCG mice were injected intra-tibially with 250,000 ATL-PDX RB in the right tibia and trabecular bone volume fraction (BV/TV) was examined by viva CT every 2 weeks (n=5). Percent change from baseline (prior to inoculation) is indicated. b) Genomic DNA PCR for amount of human cells in bone marrow of injected tibiae after 8 weeks of injection for HTLV/T lines, together with RB after 6 weeks of injection, and mouse bone marrow as negative control. c) NCG mice were injected intra-tibially with 250,000 HTLV/T lines C7 (gray), CJ2 (black) and C8 (green). Control mice were injected with PBS (open circles) or were uninjected (closed circles). BV/TV of the injected leg was determined by viva-CT before and 4- and 8-weeks post injection. Data represented as %BV/TV change from baseline (n=11-18 mice per group). d) Representative micro-CT images from mice in c. Scale bar 100μm. a, c; Linear mixed-effects model with Tukey’s method for multiplicity adjustment, ^**^ p<0.01, ^***^ p< 0.001, or ^****^ p< 0.0001 vs. Control; # p<0.05 or ## p<0.01 vs. C7.

To develop an alternative model for studying interactions between HTLV-1 infected T cells and bone cells, we generated 9 distinct HTLV-1 immortalized T cell lines (HTLV/T) from 2 different healthy donors. These HTLV/T cell lines were created by co-culturing lethally irradiated HTLV-1 producer cells with freshly isolated human PBMCs, which after 12-14 weeks established HTLV-1 infected, IL-2 dependent immortalized T cell lines. We chose three HTLV/T cell lines (C7, CJ2, C8) which grew well *in vitro*, and injected these into non-humanized 6-week-old NCG mice, intratibially. Using quantitative PCR on genomic DNA, we found that very small percentages of HTLV/T survived in mice, compared to the complete marrow replacement with ATL-PDX RB (Figure 1b). Using larger cohorts of mice, we assessed trabecular bone volume fraction (BV/TV) by vivaCT at 0, 4 and 8 weeks after inoculation. As expected, bone density increased in control mice as they are still growing at 6 weeks of age. Of the 3 HTLV/T lines implanted, CJ2 and C8 caused a profound decrease in bone mass. In contrast, even though C7-injected mice have a trend towards blunting of bone accrual, the change is not significantly different from control, and bone mass remained significantly higher than those inoculated with CJ2 or C8 (Figure 1c, d). This suggests that HTLV-1 infected T cells can drive bone loss independent of leukemia, and in fact do so even when present in very small numbers.

### 3.2 sEV from HTLV/T cell lines stimulate in vitro osteoclastogenesis

We next tested whether osteolytic HTLV/T cell line supernatants have a direct effect on osteoclasts. To this end, we cultured mouse bone marrow macrophages (mBMMs) with HTLV/T cell line supernatant (10%) in the presence of a low dose of RANKL, which is unable to form mature osteoclasts. We observed that C8 and CJ2, along with 3 other culture supernatants, had a stimulatory effect on generation of multinuclear OCs, while other cell lines had little or no effect (Figure 2a). Indeed, a similar pattern of activity among supernatants was observed in human osteoclast cultures, in which there was a significant effect on osteoclast size, greater in magnitude than the effect on their number (Supplementary Figure 1). As expected, supernatants from ATL-PDX, derived from hypercalcemic patients, consistently stimulated multinucleated osteoclast formation from murine precursors (Supplementary Figure 2a, b).

**Figure 2.**
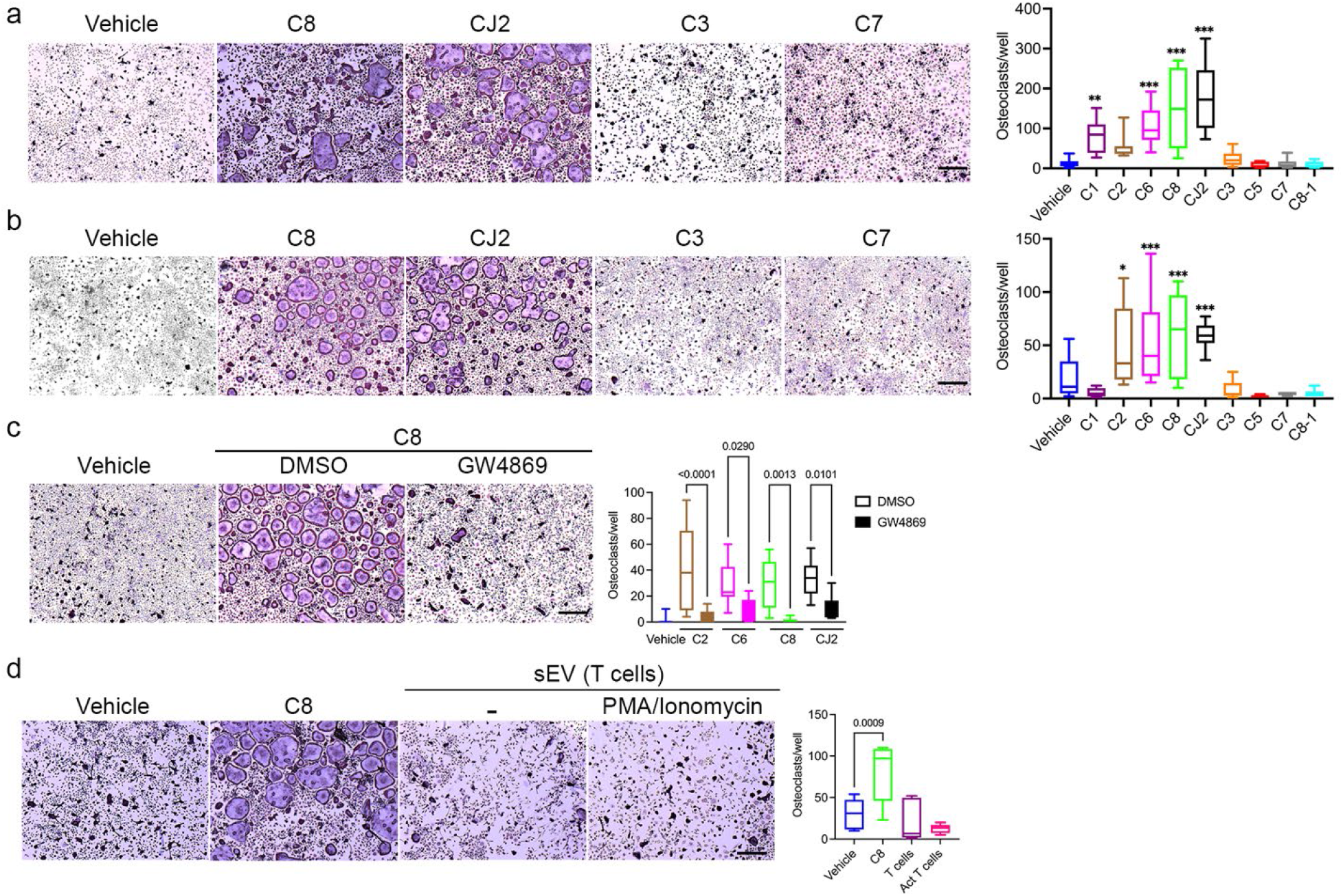
sEV from osteolytic HTLV/T cell lines stimulate *in vitro* osteoclastogenesis. a) mBMM were treated with either vehicle or 10% supernatant together with low dose RANKL for 4d. TRAP positive multinuclear cells were counted as osteoclasts. n=9, from at least 3 independent experiments. b) mBMMs were treated with ∼ 6 × 10^6^ sEV isolated from culture supernatants or vehicle for 4d. TRAP positive multinuclear cells were counted as osteoclasts. Data from 3 independent experiments, n=9. c) HTLV/T cells were pre-treated with GW4869 (10 uM) or DMSO for 5d and 10% supernatant was added during 4d of differentiation. Data from 3 biological repeats, n=9. d) sEV were isolated from IL-2 expanded human PBMC, stimulated (or not) with PMA (50ng/ml) and ionomycin (1nM), and added to osteoclastognic cultures for 4d. Results from 2 independent experiments, n=6. Scale bars 500 μm. All statistics one way ANOVA, ^*^p<0.05, ^**^p<0.01, ^***^p<0.001.

We previously showed that ATL cells express RANKL, driven by viral HBZ expression, and anti-RANKL antibody treatment offered moderate protection from bone loss in HTLV-1-infected humanized mice^6^. Therefore, we quantitated the cellular mRNA level of RANKL, the RANKL antagonist OPG, and their ratio in HTLV/T clones. They expressed variable amounts of these factors and displayed no correlation to OC numbers (Supplementary Figure 2c). We also observed that ATL-PDX display varying expression of RANKL and OPG (Supplementary Figure 2d). This suggested that additional factors in the supernatant other than RANKL and OPG might dictate the osteolytic phenotype.

Because sEV have been shown to play a role in HTLV-1 transmission and progression in a humanized mouse model^24^, we hypothesized that the sEV carry the osteoclast-stimulatory activity of HTLV/T supernatants. We therefore precipitated sEV from culture supernatant using commercial kits, and confirmed their size distribution and concentration (Supplementary Figure 3a). We also demonstrated their uptake by mBMM (Supplementary Figure 3b). The sEV isolated from stimulatory supernatants similarly promoted multinucleated OC formation, whether derived from ATL-PDX or HTLV/T lines (Supplementary Figure 4 and Figure 2b), except for sEV from HTLV/T C1 supernatant. To further strengthen the evidence for a stimulatory role of sEV rather than soluble factors in culture supernatants, we pre-treated HTLV/T cell lines with a chemical inhibitor of EV generation (GW4869), which did not affect cell viability (Supplementary Figure 5), and tested the activity of the resulting supernatants. In all cases, the number of mature osteoclasts were decreased when these EV-depleted supernatants were added to mBMM, compared to supernatants from DMSO control treated cells (Figure 2c).

HTLV-1 infection leads to T cell activation^33^. Since activated T cells are known to stimulate osteoclastogenesis, we sought to determine whether such activation, rather than viral infection itself, leads to production of osteoclast-active sEV. We utilized hPBMC, cultured in IL-2 with or without stimulation by PMA and ionomycin for 2 days, and isolated sEV. We observed that sEV from activated T cell cultures failed to impact osteoclasts (Figure 2d). These results indicate that the osteoclastogenic activity of sEV is specific for those derived from T cells infected by HTLV-1.

### 3.3 HTLV-1 infected T cells promote the maturation of osteoclasts and do not act on precursor bone marrow macrophages

To determine if HTLV/T stimulated the OC differentiation program, we examined osteoclast-specific gene expression after treatment with culture supernatants. Surprisingly, we found no increase in marker levels compared to vehicle controls (Supplementary Figure 6a). This prompted us to examine which stage of osteoclastogenesis was targeted by HTLV/T cells. Osteoclast differentiation can be broadly divided into two stages, generation of committed pre-osteoclasts and their subsequent fusion to multinucleated giant cells^35^. We performed a time course study where we added supernatant either for the first two days or last two days of differentiation, which indicated that HTLV/T targeted the latter maturation stage (Figure 3a). Similarly, pre-osteoclasts generated in low dose RANKL for 2 days and then treated with sEV for an additional day formed more multinucleated osteoclasts than those treated with PBS (Figure 3b). As with supernatant-treated cells, there was no change in expression of differentiation genes in sEV-treated cultures (Supplementary Figure 6b). To determine if the generated osteoclasts were functional, we cultured pre-osteoclasts as above, plated them on bone slices, then added supernatant for 5 more days, and examined resorption activity. As expected, HTLV/T supernatants with high activity on multinucleation also had greater effects on resorptive potential (Figure 3c). This suggests that HTLV/T culture supernatants enhanced the fusion of pre-osteoclasts to generate mature, bone-resorbing cells.

**Figure 3.**
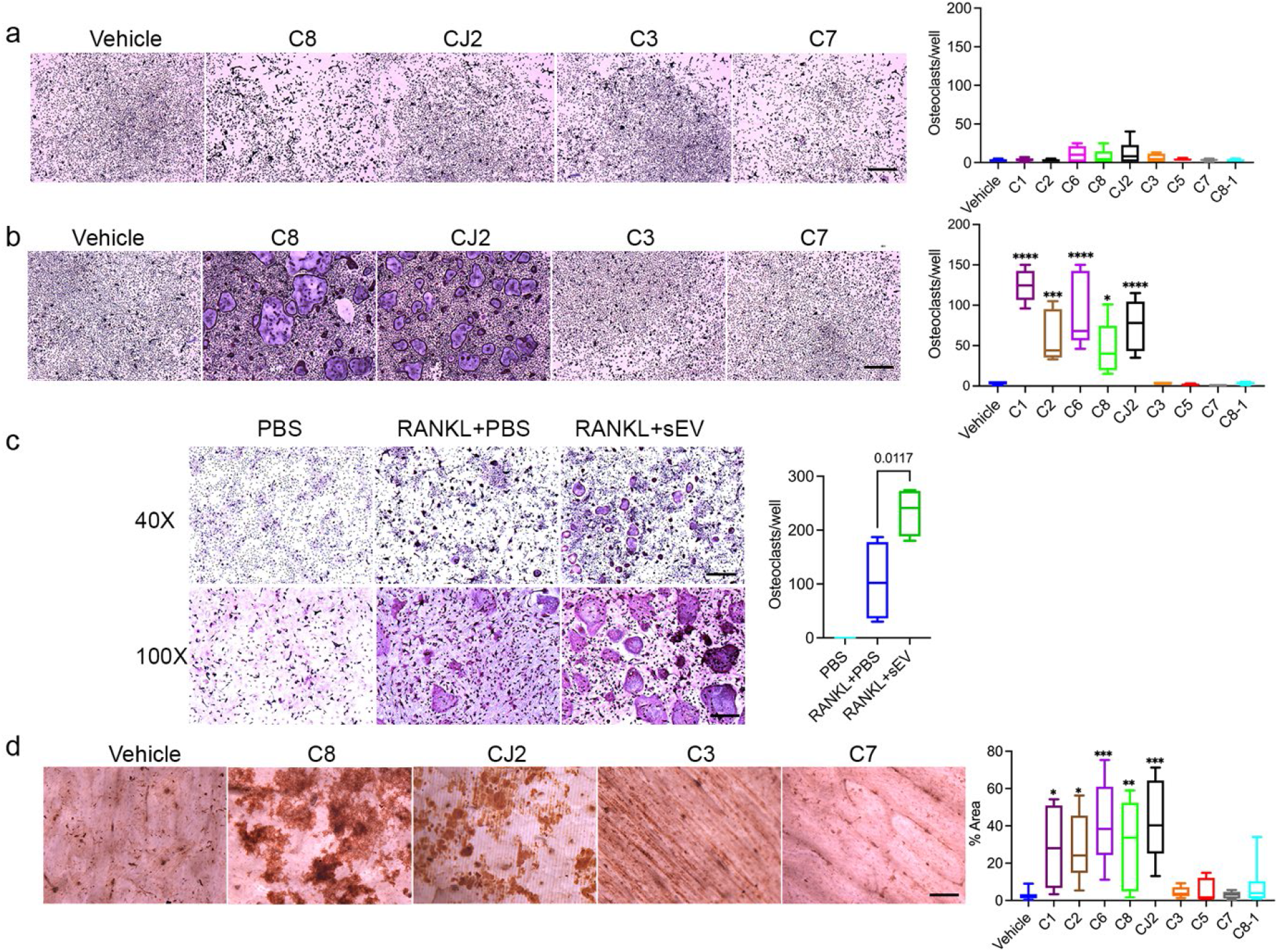
HTLV/T cells modulate maturation of osteoclasts. a,b) Supernatants were added to osteoclastogenic mBMM cultures during first 2d (a) or last 2d (b). Cells were fixed and TRAP stained after 4d, and osteoclasts were counted. c) Pre-osteoclasts were generated by treating mBMM with 50 ng/ml of RANKL for 2d. These were then cultured with low dose RANKL with or without sEV (from HTLV/T C8) for an additional 24 h. c) Pre-osteoclasts were lifted and replated on bone slices, treated with supernatant for 5d, and resorption pits were stained. Resorption area was measured using ImageJ. n=6-9, from 2-3 independent experiments. Scale bars: 500μm - a, b, and c (upper panel); 200μm - c (lower panel), d. All statistics one way ANOVA, ^*^ p<0.05, ^**^p<0.01, ^***^p<0.001, ^****^p<0.0001.

### 3.4 Profiling of sEV from HTLV-1 infected T cells identifies proteins and miRNAs that regulate osteoclastogenesis

We performed transmission electron microscopic (TEM) analysis of both HTLV/T and ATL-PDX sEV, and found them to be electro-lucent and lacking dense virion cores, confirming the absence viral particles in our sEV preparations (Figure 4a, Supplementary Figure 7a), similar to what others have demonstrated with ultracentrifugation-based isolation of sEV from HTLV-1 infected cells^22^. We then performed LC-MS/MS on osteoclast-active (n=4) and inactive (n=3) HTLV/T sEV groups. Analysis confirmed the presence of sEV markers according to Minimal Information for Studies of Extracellular Vesicles (MISEV) guidelines^36^ in all 7 sEV preparations (Supplementary Figure 7b). HTLV/T sEV carry only 3 HTLV-1 viral polypeptides, of which two (Env and Tax) are only present in sEV with stimulatory effects on osteoclasts (Figure 4b). Notably, neither RANKL nor OPG was detected in any of the sEV. To gain further insight into components of the sEV, we examined protein abundance by heat map (Figure 4c) and clustering between the groups by PCA plot (Figure 4d). These results both showed separation of active versus inactive groups of sEV. Furthermore, correlation analysis revealed that among 5248 proteins carried in the cargo, 968 (18.5%) proteins were highly correlated with the osteoclastogenic phenotype (Figure 4e). Of those highly correlated proteins, gene ontology analysis showed that most of the proteins are cytoskeletal, GTPases, and kinases, which could have a role in influencing fusion and resorptive function (Figure 4f). We used STRING to understand the protein-protein interactions and found that osteoclast differentiation and TCR signaling pathways overlapped in this analysis (Supplementary Figure 8). The osteoclast differentiation proteins were enriched in the active group, (Figure 4g) while inactive sEV had an abundance of thrombospondins (Figure 4h). We also performed miRNA sequencing on the active sEV group. As expected, these sEV were rich in miRNAs and, the top miRNAs in all 4 active sEV preparations were similar (Supplementary Figure 9). Interestingly, of those, three miRNAs (mir155, mir21, mir92a1) were previously reported to have a positive effect on osteoclast differentiation^37,38,39^. These results suggest that both the protein and miRNAs in sEV cargo have the potential to affect OC maturation and function.

**Figure 4.**
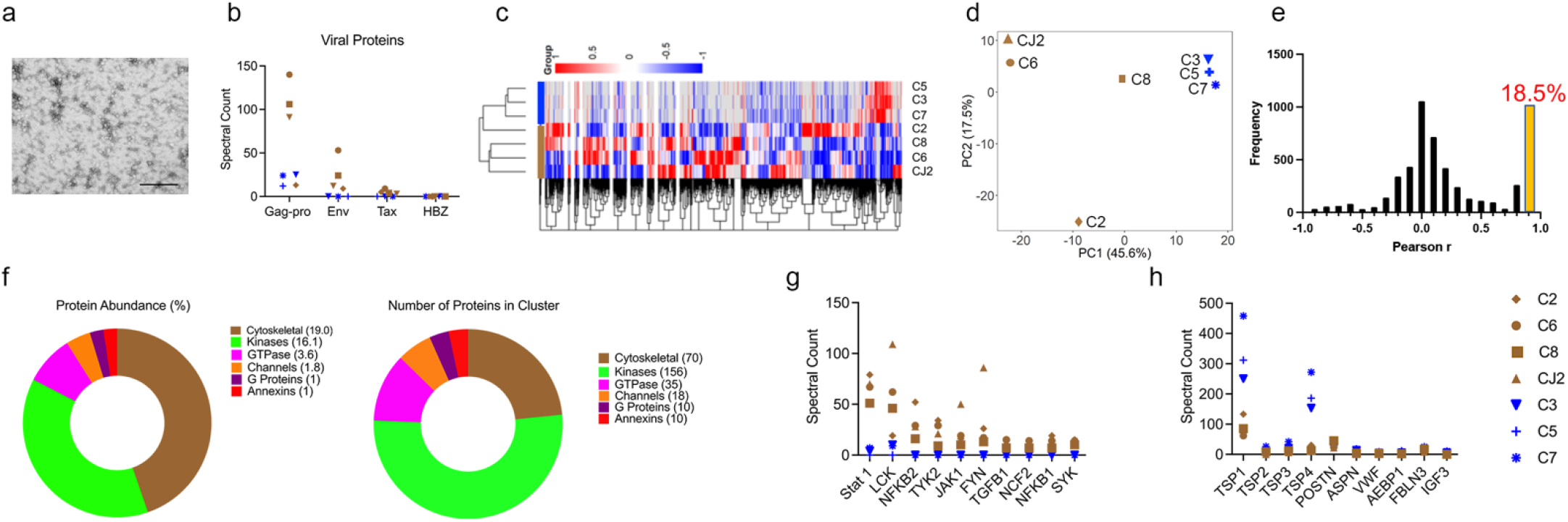
Proteomics of sEV suggests multiple possible candidates modulating osteoclastogenesis. a) Representative transmission electron micrograph, of sEV isolated from C8. Scale bar: 100 nm b-h) LC-MS/MS analysis was performed on proteins isolated from sEV with positive (n=4, brown), and negligible (n=3, blue) effect on osteoclast differentiation. b) Analysis of viral proteins shows presence of gag-pro in all sEV, but Env (gp62) and Tax only in sEV that stimulate osteoclasts. HBZ was not detected in any. c-h) Analysis of cellular proteins, c) Heat map, d) PCA plot, e) Pearson’s correlation, f) Gene ontogeny analysis. g) STRING network analysis displaying enriched proteins in sEV with high osteoclastogenic effect, h) STRING network analysis displaying enriched proteins in sEV without osteoclastogenic effect.

### 3.5 sEV from HTLV-1 infected T cells are sufficient to cause in vivo osteolysis in non-tumor bearing mice

We next sought to establish whether HTLV/T-derived sEV directly induce osteolysis *in vivo*. As we had determined that sEV induced formation of mature osteoclasts only in RANKL-primed cultures, we used a similar approach in a mouse osteolysis model. We injected RANKL over the calvarium for 2 days to initiate OC formation, then treated with a low dose of RANKL with or without sEV from C8, a stimulatory HTLV/T line, for 3 more days. Mice were euthanized 2 days after the last treatment, allowing time for osteoclasts to resorb bone. Consistent with its effect *in vitro*, application of C8 sEV to calvaria increased pit formation and apparent width of sutures, leading to a marked decrease in bone mass, compared to those treated with RANKL alone (Figure 5a). Upon histological examination, we observed a significant increase in TRAP positive osteoclasts (Figure 5b). These results indicate that sEV released by HLTV-1 infected cells have the ability to stimulate osteoclasts *in vivo*, leading to osteolysis.

**Figure 5.**
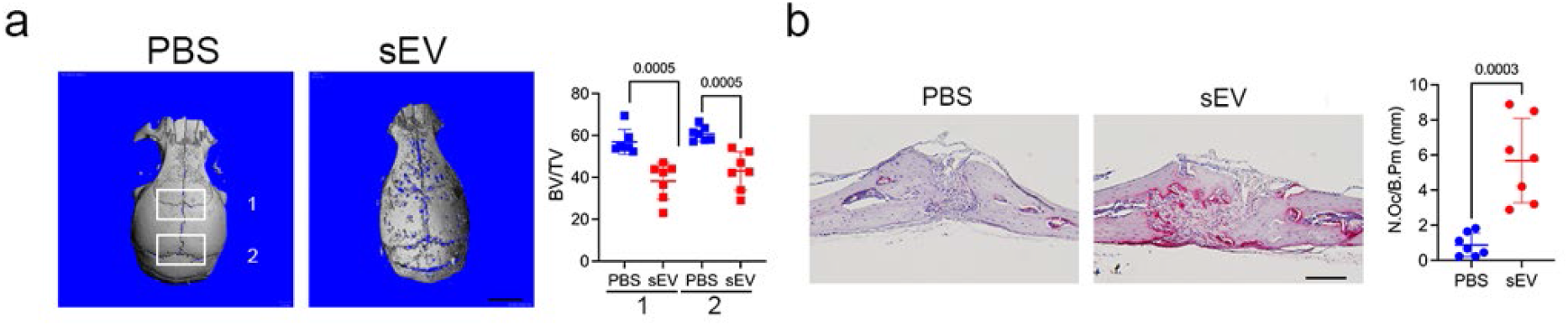
HTLV/T sEVs cause bone loss in calvarial osteolysis model. Mice were injected with RANKL together with PBS or sEV (9×10^7^) from HTLV/T C8 subcutaneously over the calvaria. a) Calvaria were scanned using micro-CT and bone volume fraction (BV/TV) was determined in 2 regions, as indicated. Scale bar: 1 mm. b) Calvaria were decalcified, and paraffin embedded sections were subjected to TRAP staining. Osteoclast number per bone perimeter (N.Oc/B.Pm) were counted. n=7/group. Scale bar: 100 μm. Unpaired t-test, p values indicated.

## 4 Discussion

To understand the role of HTLV-1 infection in bone loss, we generated HTLV-1 infected, immortalized T cell lines and implanted them intra-tibially. Interestingly, although these cells persisted at low levels in the bone marrow over 8 weeks, some HTLV/T cell lines caused progressive bone loss. This osteolytic activity correlated with *in vitro* ability of supernatants derived from these cells to stimulate the formation of mature osteoclasts. The finding is quite remarkable given that very few HTLV/T cells survive in the bone marrow, but they have a strong and sustained effect, causing a similar amount of bone loss to ATL-PDX models which replace the normal marrow constituents. Further analysis indicated that sEV, rather than RANKL or other soluble proteins, were the primary drivers of bone loss. Consistent with their *in vitro* effect, isolated sEV injected subcutaneously caused rapid osteolysis. Although activated T cells are known to stimulate osteoclastogenesis via RANKL expression, their sEV did not, indicating that the osteolytic potential of sEV was specific to HTLV-1 infection.

Expression of sEV markers, defined by MISEV guidelines^36^, were similar between the sEV from the HTLV/T cell lines with or without effects on osteoclast differentiation. Similar to sEV from HTLV-1-infected cells isolated by other groups, these sEV lacked complete viral particles, and were also devoid of RANKL and OPG. Our osteoclastogenic sEV shared other features with prior studies. Jaworski et al, 2014 found that HTLV-1 infected T cells release sEV with viral protein Tax^22^. Interestingly, in our study, Env and Tax were only present in sEV with osteoclast stimulatory activity. Env in particular has been found to be fusogenic^40^ and may play a similar role to enhance multinucleation of osteoclasts in our cultures utilizing suboptimal doses of RANK. Formation of very large, hypermultinucleated osteoclasts is characteristic in Paget’s Disease, which can be associated with paramyxovirus infection^41^. Notably, however, we have not observed sEV-induced multinucleation beyond the levels obtained with higher, optimized doses of RANKL in the absence of sEV. Osteoclastogenic sEV were also enriched for many proteins identified in other studies to be involved in osteoclast differentiation and function^35^. Additionally, the most abundant miRNAs, also found in sEV by El-Saghir^28^, were miRNA155 and miRNA21, both with osteoclastogenic effects^37,38^. Thus, the osteoclast-active sEV bear a complex cargo with multiple components with the potential to enhance osteoclast maturation and resorptive function. It is therefore unlikely that modulation of a single component will eliminate their activity.

The ability of very small numbers of HTLV/T cells to cause bone loss in immunodeficient mice *in vivo* was surprising. These cells are dependent on human IL-2 *in vitro*, and we did not transplant human hematopoietic stem cells (i.e. humanization) nor did we provide an exogenous source of IL-2. It is possible that provision of IL-2 or other T cell supportive cytokines would lead to significant lymphoproliferation *in vivo*. Nevertheless, this fortuitous finding of extensive osteolysis points to the presence of a potent mediator. Important to our conclusion that sEV are the primary candidates of osteoclast activation is the absence of an *in vivo* effect of implanted C7 HTLV/T cells, which persist at similar levels to highly osteolytic CJ2.

Neither the whole supernatant nor isolated sEV from C7 are stimulatory *in vitro*. Nano Sight analysis confirmed similar concentrations and size distributions of isolated sEV, and proteomic analysis showed that C7, and other inactive sEV, were enriched in some proteins compared to the active sEV, suggesting a distinct cargo rather than defective sEV production by these cell lines. We thought it was important to directly apply sEV *in vivo* to assess their osteolytic effects. Because the HTLV/T cells grow slowly, we were unable to inject large numbers systemically over weeks, as others have done for solid tumor-derived sEV^25,26^. Instead we used only 3 injections locally over the calvarium, and the number of sEV was 1000-fold lower than these other studies. Whether HTLV/T-derived sEV are particularly potent in their effects on bone will require direct comparisons with sEV from other sources. Additionally, the reciprocal role of the bone microenvironment on HTLV-1 infected T cells, prior to and after transformation, remains to be investigated.

Interestingly, we observed that sEV from HTLV/T cells with minimal effect in osteoclastogenesis were enriched in thrombospondins, markers associated with asymptomatic carriers of HTLV-1^42^. It is intriguing to speculate that a preponderance of sEV with thrombospondins might predict a lower propensity to ATL progression, compared to those with enrichment of markers such as Env found at high levels in osteolytic sEV. A previous proteomic analysis of plasma-derived EVs from HTLV-1 infected patients failed to detect viral proteins^43^. Thus, isolation of sEV specifically from T cells might be needed to achieve adequate sensitivity. Significantly, more investigation of potential prognostic utility is warranted of sEV from HTLV-1 infected patients with a range of disease types, including asymptomatic, and with HTLV-associated myelopathy, leukemia and lymphoma.

In sum, our data shows that HTLV-1 infection of T cells leads to production of sEV that can drive bone loss *in vivo*, independent of the development of systemic leukemia. Our understanding of sEV as a modulator of osteoclastogenesis could be beneficial for providing a better understanding of bone-tumor interactions, designing new clinical therapies, and using sEV for predicting disease progression.

## Author Contributions

**Nitin K Pokhrel**: Investigation (lead); methodology (lead); formal analysis (lead); writing – original draft (lead); writing – review and editing (lead). **Amanda Panfil**: conceptualization (supporting), methodology (supporting), writing – review and editing (supporting). **Ancy Joseph**: methodology and investigation (supporting). **Haniya Habib, Sham Seeniraj**: formal analysis (supporting). **Daniel Rauch, Qiang Zhang, Rajeev Aurora, William Park**: formal analysis (supporting). **Linda Cox, Robert Sprung, Petra Gilmore**: investigation (supporting). **Reid Townsend**: conceptualization (supporting), methodology (supporting), supervision. **Lee Ratner, Katherine N Weilbaecher**: conceptualization (supporting), formal analysis (supporting), review and editing (supporting), funding acquisition, supervision. **Deborah J Veis**: conceptualization (lead), writing – original draft (lead), writing – review and editing (lead), funding acquisition, project administration, administration, supervision.

## Acknowledgements

We thank Crystal Idleburg and Samantha Coleman Cathcart in the Washington University Musculoskeletal Histology and Morphometry Core for assistance with histology and Michael Brodt in the Structure and Strength Core for microCT analysis.

## References

1. Mehta-Shah N, Ratner L, Horwitz SM. Adult T-Cell Leukemia/Lymphoma. J Oncol Pract. 2017 Aug;13(8):487–492. doi: 10.1200/JOP.2017.021907. PMID: 28796966; PMCID: PMC6366298.

2. Kiyokawa T, Yamaguchi K, Takeya M, Takahashi K, Watanabe T, Matsumoto T, et al. Hypercalcemia and osteoclast proliferation in adult T-cell leukemia. Cancer. 1987 Mar 15;59(6):1187–91.

3. Prager D, Rosenblatt JD, Ejima E. Hypercalcemia, parathyroid hormone-related protein expression and human T-cell leukemia virus infection. Leuk Lymphoma. 1994 Aug;14(5-6):395–400.

4. Kohart NA, Elshafae SM, Supsahvad W, Alasonyalilar-Demirer A, Panfil AR, Xiang J, et al. Mouse model recapitulates the phenotypic heterogeneity of human adult T-cell leukemia/lymphoma in bone. J Bone Oncol. 2019 Aug 20;19:100257.

5. Shu ST, Martin CK, Thudi NK, Dirksen WP, Rosol TJ. Osteolytic bone resorption in adult T-cell leukemia/lymphoma. Leuk Lymphoma. 2010 Apr;51(4):702–14.

6. Xiang J, Rauch DA, Huey DD, Panfil AR, Cheng X, Esser AK, et al. HTLV-1 viral oncogene HBZ drives bone destruction in adult T cell leukemia. JCI Insight. 2019 Oct 3;4(19).

7. Nosaka K, Miyamoto T, Sakai T, Mitsuya H, Suda T, Matsuoka M. Mechanism of hypercalcemia in adult T-cell leukemia: overexpression of receptor activator of nuclear factor kappaB ligand on adult T-cell leukemia cells. Blood. 2002 Jan 15;99(2):634–40.

8. Okada Y, Tsukada J, Nakano K, Tonai S, Mine S, Tanaka Y. Macrophage inflammatory protein-1alpha induces hypercalcemia in adult T-cell leukemia. J Bone Miner Res. 2004 Jul;19(7):1105–11.

9. Parrula C, Zimmerman B, Nadella P, Shu S, Rosol T, Fernandez S, et al. Expression of tumor invasion factors determines systemic engraftment and induction of humoral hypercalcemia in a mouse model of adult T-cell leukemia. Vet Pathol. 2009 Sep;46(5):1003–14.

10. Grunbaum A, Kremer R. Parathyroid hormone-related protein (PTHrP) and malignancy. Vitam Horm. 2022;120:133–177.

11. trivedi T, Pagnotti GM, Guise TA, Mohammad KS. The Role of TGF-β in Bone Metastases. Biomolecules. 2021 Nov 6;11(11):1643.

12. Rieunier G, Wu X, Macaulay VM, Lee AV, Weyer-Czernilofsky U, Bogenrieder T. Bad to the Bone: The Role of the Insulin-Like Growth Factor Axis in Osseous Metastasis. Clin Cancer Res. 2019 Jun 15;25(12):3479–3485.

13. taipaleenmäki H. Secreted microRNAs in bone metastasis. J Bone Miner Metab. 2023 May;41(3):358–364.

14. Foster BM, Shi L, Harris KS, Patel C, Surratt VE, Langsten KL, et al. Bone Marrow-Derived Stem Cell Factor Regulates Prostate Cancer-Induced Shifts in Pre-Metastatic Niche Composition. Front Oncol. 2022 Apr 19;12:855188.

15. Kode A, Manavalan JS, Mosialou I, Bhagat G, Rathinam CV, Luo N, et al. Leukaemogenesis induced by an activating β-catenin mutation in osteoblasts. Nature. 2014 Feb 13;506(7487):240–4.

16. Giannakoulas N, Ntanasis-Stathopoulos I, Terpos E. The Role of Marrow Microenvironment in the Growth and Development of Malignant Plasma Cells in Multiple Myeloma. Int J Mol Sci. 2021 Apr 24;22(9):4462.

17. Niewiesk S. Animals Models of Human T Cell Leukemia Virus Type I Leukemogenesis. ILAR J. 2016;57(1):3–11.

18. Gao L, Deng H, Zhao H, Hirbe A, Harding J, Ratner L, et al. HTLV-1 Tax transgenic mice develop spontaneous osteolytic bone metastases prevented by osteoclast inhibition. Blood. 2005 Dec 15;106(13):4294–302.

19. Esser AK, Rauch DA, Xiang J, Harding JC, Kohart NA, Ross MH, et al. HTLV-1 viral oncogene HBZ induces osteolytic bone disease in transgenic mice. Oncotarget. 2017 Aug 27;8(41):69250–69263.

20. Urabe F, Patil K, Ramm GA, Ochiya T, Soekmadji C. Extracellular vesicles in the development of organ-specific metastasis. J Extracell Vesicles. 2021 Jul;10(9):e12125.

21. Becker A, Thakur BK, Weiss JM, Kim HS, Peinado H, Lyden D. Extracellular Vesicles in Cancer: Cell-to-Cell Mediators of Metastasis. Cancer Cell. 2016 Dec 12;30(6):836–848.

22. Jaworski E, Narayanan A, Van Duyne R, Shabbeer-Meyering S, Iordanskiy S, Saifuddin M, et al. Human T-lymphotropic virus type 1-infected cells secrete exosomes that contain Tax protein. J Biol Chem. 2014 Aug 8;289(32):22284–305.

23. Pinto DO, DeMarino C, Pleet ML, Cowen M, Branscome H, Al Sharif S, et al. HTLV-1 Extracellular Vesicles Promote Cell-to-Cell Contact. Front Microbiol. 2019 Sep 18;10:2147.

24. Pinto DO, Al Sharif S, Mensah G, Cowen M, Khatkar P, Erickson J, et al. Extracellular vesicles from HTLV-1 infected cells modulate target cells and viral spread. Retrovirology. 2021 Feb 23;18(1):6.

25. Wu K, Feng J, Lyu F, Xing F, Sharma S, Liu Y, et al. Exosomal miR-19a and IBSP cooperate to induce osteolytic bone metastasis of estrogen receptor-positive breast cancer. Nat Commun. 2021 Aug 31;12(1):5196.

26. Urabe F, Kosaka N, Yamamoto Y, Ito K, Otsuka K, Soekmadji C, et al. Metastatic prostate cancer-derived extracellular vesicles facilitate osteoclastogenesis by transferring the CDCP1 protein. J Extracell Vesicles. 2023 Mar;12(3):e12312.

27. Pitari MR, Rossi M, Amodio N, Botta C, Morelli E, Federico C, et al. Inhibition of miR-21 restores RANKL/OPG ratio in multiple myeloma-derived bone marrow stromal cells and impairs the resorbing activity of mature osteoclasts. Oncotarget. 2015 Sep 29;6(29):27343–58.

28. El-Saghir J, Nassar F, Tawil N, El-Sabban M. ATL-derived exosomes modulate mesenchymal stem cells: potential role in leukemia progression. Retrovirology. 2016 Oct 19;13(1):73.

29. Maksimova V, Wilkie T, Smith S, Phelps C, Melvin C, Yu L, et al. HTLV-1 Hbz protein, but not hbz mRNA secondary structure, is critical for viral persistence and disease development. PLoS Pathog. 2023 Jun 16;19(6):e1011459.

30. Maksimova V, Smith S, Seth J, Phelps C, Niewiesk S, Satou Y, et al. HTLV-1 intragenic viral enhancer influences immortalization phenotype in vitro, but is dispensable for persistence and disease development in animal models. Front Immunol. 2022 Jul 25;13:954077.

31. Novack DV, Yin L, Hagen-Stapleton A, Schreiber RD, Goeddel DV, Ross FP, et al. The IkappaB function of NF-kappaB2 p100 controls stimulated osteoclastogenesis. J Exp Med. 2003 Sep 1;198(5):771–81.

32. takeshita S, Kaji K, Kudo A. Identification and characterization of the new osteoclast progenitor with macrophage phenotypes being able to differentiate into mature osteoclasts. J Bone Miner Res. 2000 Aug;15(8):1477–88.

33. Kataoka K, Nagata Y, Kitanaka A, Shiraishi Y, Shimamura T, Yasunaga J, et al. Integrated molecular analysis of adult T cell leukemia/lymphoma. Nat Genet. 2015 Nov;47(11):1304–15.

34. Veis DJ, O’Brien CA. Osteoclasts, Master Sculptors of Bone. Annu Rev Pathol. 2023 Jan 24;18:257–281. doi: 10.1146/annurev-pathmechdis-031521-040919. Epub 2022 Oct 7. PMID: 36207010.

35. théry C, Witwer KW, Aikawa E, Alcaraz MJ, Anderson JD, Andriantsitohaina R, et al. Minimal information for studies of extracellular vesicles 2018 (MISEV2018): a position statement of the International Society for Extracellular Vesicles and update of the MISEV2014 guidelines. J Extracell Vesicles. 2018 Nov 23;7(1):1535750.

36. Mao Z, Zhu Y, Hao W, Chu C, Su H. MicroRNA-155 inhibition up-regulates LEPR to inhibit osteoclast activation and bone resorption via activation of AMPK in alendronate-treated osteoporotic mice. IUBMB Life. 2019 Dec;71(12):1916–1928.

37. Hu CH, Sui BD, D. FY, Shuai Y, Zheng CX, Zhao P, et al. miR-21 deficiency inhibits osteoclast function and prevents bone loss in mice. Sci Rep. 2017 Feb 27;7:43191. Erratum in: Sci Rep. 2022 Dec 14;12(1):21627.

38. Yu L, Sui B, Fan W, Lei L, Zhou L, Yang L, et al. Exosomes derived from osteogenic tumor activate osteoclast differentiation and concurrently inhibit osteogenesis by transferring COL1A1-targeting miRNA-92a-1-5p. J Extracell Vesicles. 2021 Jan;10(3):e12056.

39. Hoshino H. Cellular Factors Involved in HTLV-1 Entry and Pathogenicity. Front Microbiol. 2012 Jun 21;3:222.

40. Kurihara N, Hiruma Y, Yamana K, Michou L, Rousseau C, Morissette J, et al. Contributions of the measles virus nucleocapsid gene and the SQSTM1/p62(P392L) mutation to Paget’s disease. Cell Metab. 2011 Jan 5;13(1):23–34.

41. Ferreira CES, Martins ML, Brito-Melo GEA, Carvalho LD de, Carneiro-Proiett AB de F, Namen-Lopes MS, et al. Thrombospondin-1 expression in human t-lymphotropic virus 1 asymptomatic carriers and patients with htlv-associated myelopathy/tropical spastic paraparesis. Rev Patol Trop. 2012 Oct 22;41(3).

42. Jeannin P, Chaze T, Giai Gianetto Q, Matondo M, Gout O, Gessain A, et al. Proteomic analysis of plasma extracellular vesicles reveals mitochondrial stress upon HTLV-1 infection. Sci Rep. 2018 Mar 26;8(1):5170.

