## Supplemental Data for "HTLV-1 infected T cells cause bone loss via small extracellular vesicles"

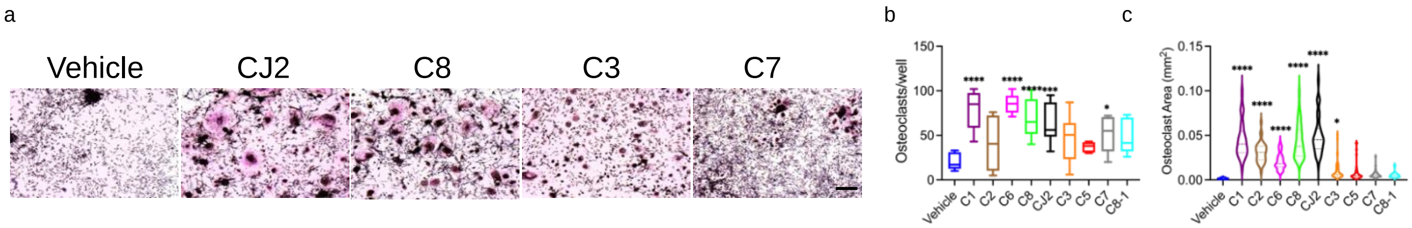


Supplementary Figure 1. Supernatants from HTLV/T cell lines variably affect differentiation of human PBMC to osteoclasts.

a) Total human PBMC were treated with 10% HTLV/T supernatant. The cells were cultured for 5-6 days, fixed and subjected to TRAP staining. b) Multinucleated osteoclasts with >3 nuclei were counted. c) Average area of the osteoclasts was measured using Image J. Data from at least two independent experiments. n=6-9. Scale bar 200 μm , One-way anova: * p<0.05; *** p<0.001; ****p<0.0001.

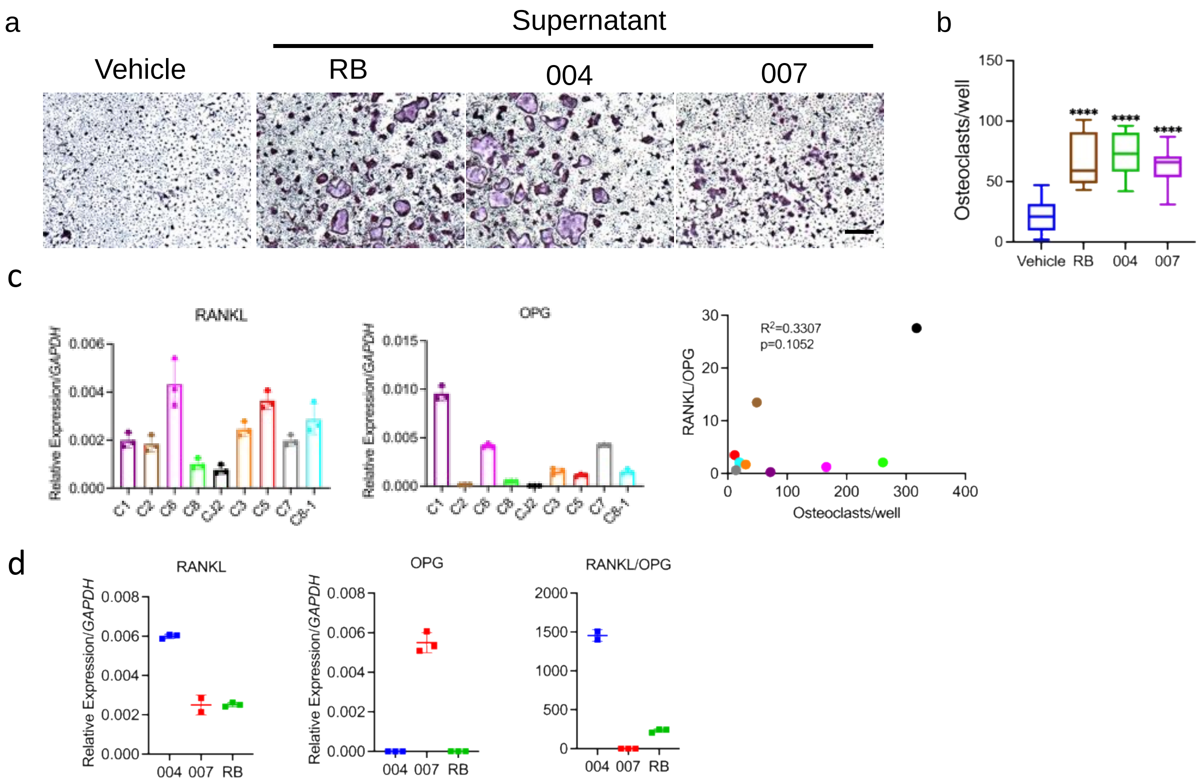


Supplementary Figure 2. Stimulation of osteoclastogenesis does not correlate with RANKL, OPG expression levels or RANKL/OPG ratio. a-b) Mouse bone marrow macropahges (mBMM) were treated with 10% supernatant from ATL-PDX. Scale bar: 500 μm, TRAP positive multinuclear cells (MNCs) were counted as osteoclasts. 3 independent experiments, n=9. One-way anova, ****, $p<0.0001$. c) Cellular mRNA expression level of RANKL and OPG by HTLV/T cell lines and their correlation to OC numbers in mBMM cultures. Pearson correlation, two-tailed, 95% CI. d) Cellular mRNA expression level of RANKL and OPG by ATL-PDX.

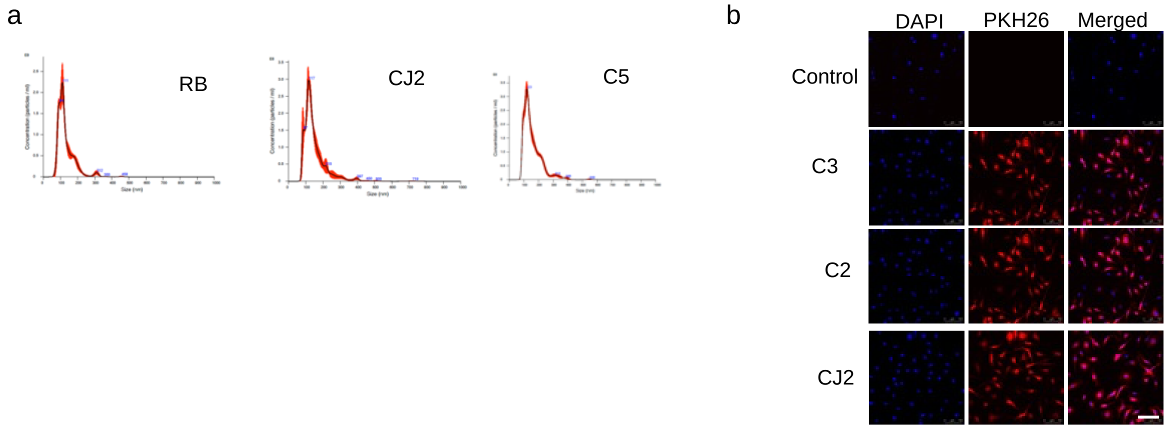


Supplementary Figure 3. sEV characterization. sEV were isolated from culture supernatants from ATL-PDX and HTLV/T. a) sEV were analyzed by Nano Sight 300 for examination of their size distribution and particle count. All had similar characteristics, including from the osteoclast-active and inactive HTLV/T groups. b) sEV were labelled with fluorescent dye PKH 26 and subsequently cultured with mBMM for 4 h. The cells were then fixed and imaged. Scale bar 100 μm.

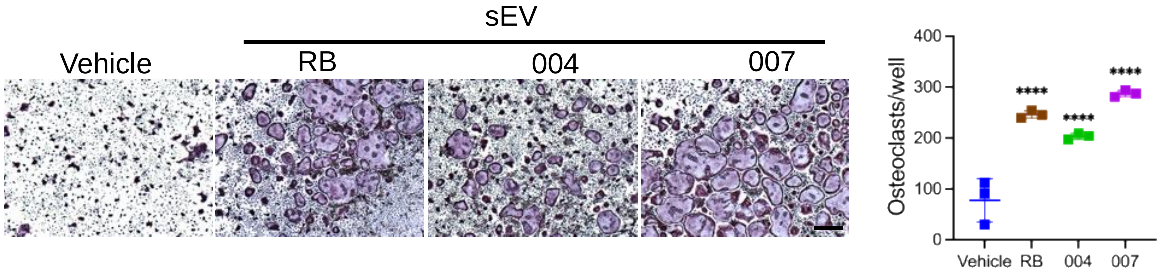


Supplementary Figure 4: ATL-PDX sEV stimulate osteoclastogenesis. sEV were isolated from culture supernatant and added to mBMM culture for 4d. TRAP positive multinuclear cells were counted as osteoclasts. Data represents technical replicates. Scale bar 500 µm, One-way anova, **** $p<0.0001$.

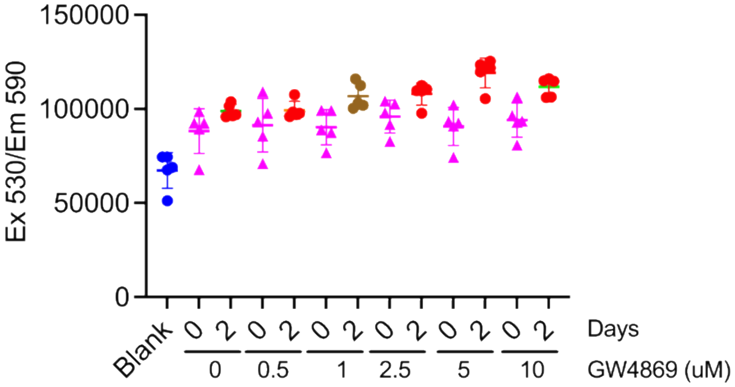


Supplementary Figure 5: Effect of GW4869 on cell viability. HLTV/T CJ2 was treated with increasing amounts of GW4869 in DMSO as indicated and cell viability is assessed using MTT assay after 2d.

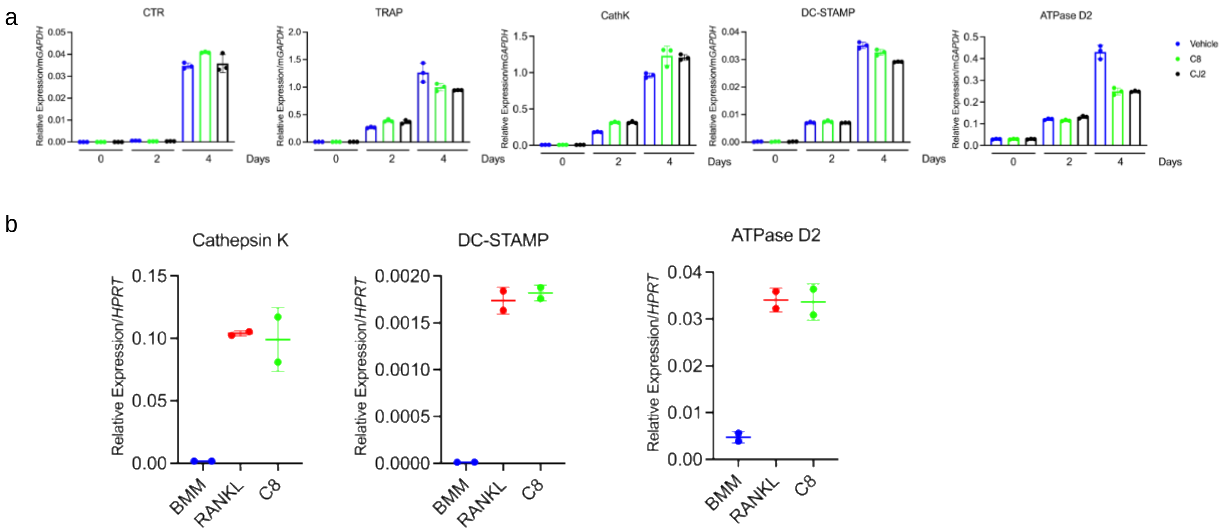


Supplementary Figure 6. sEV have no effect on expression of osteoclast marker genes. a) Cellular expression level of osteoclast differentiation genes during osteoclastogenic culture of mBMM with HTLV/T supernatant from the indicated HTLV/T lines. Data represents technical replicates. b) Pre-osteoclasts were generated from mBMM with a 2 d RANKL treatment, and sEV were added for an additional 24h prior to assessment of expression level of genes associated with osteoclast fusion. Data representative of at least 2 independent experiments.

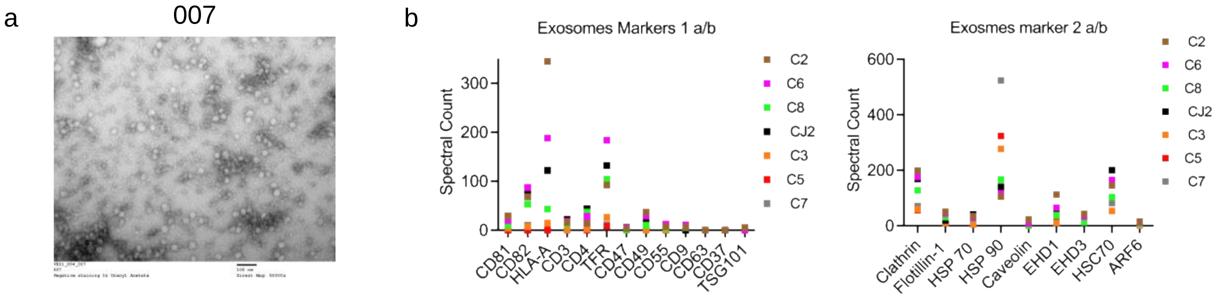


Supplementary Figure 7: sEV from HTLV/T cells do not contain virus, but do conform to MISEV guidelines for exosomes. a) sEV were negatively stained with uranyl acetate and examined by TEM. b) LC-MS/MS analysis was performed on sEV preparations. Exosomal marker distribution (spectral count) is presented, as defined by MISEV guidelines[31].

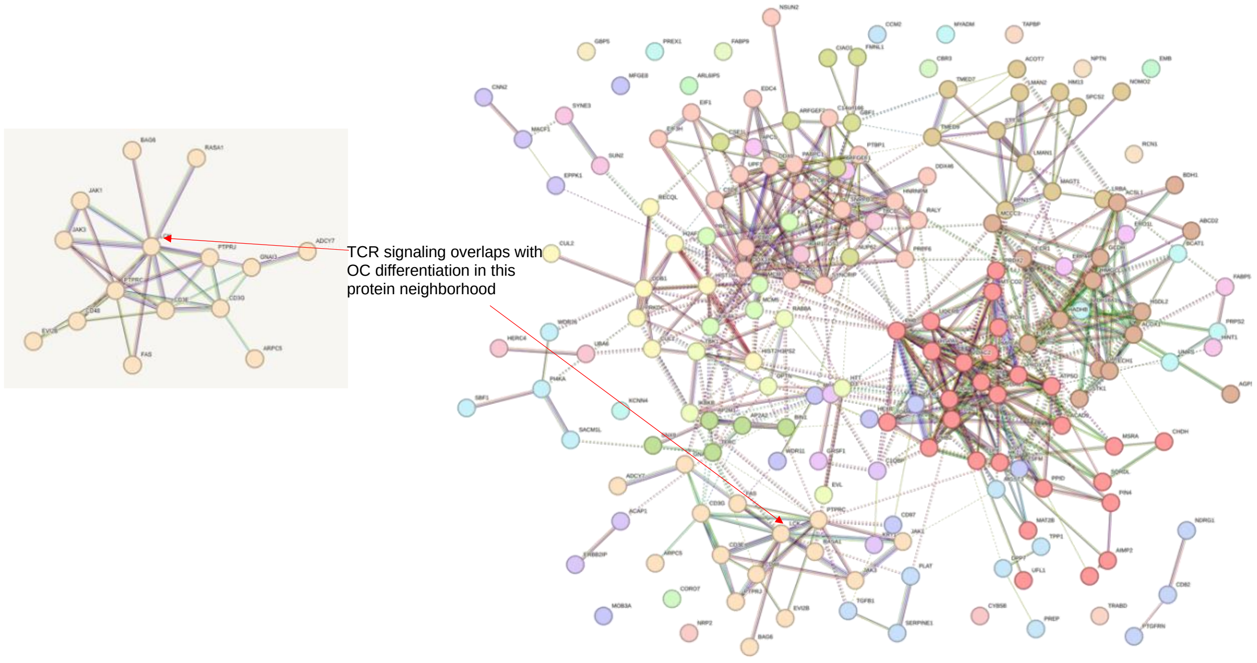


Supplementary Figure 8: Interaction of TCR signalling with osteoclast differentiation demonstrated by STRING analysis of sEV proteomics.

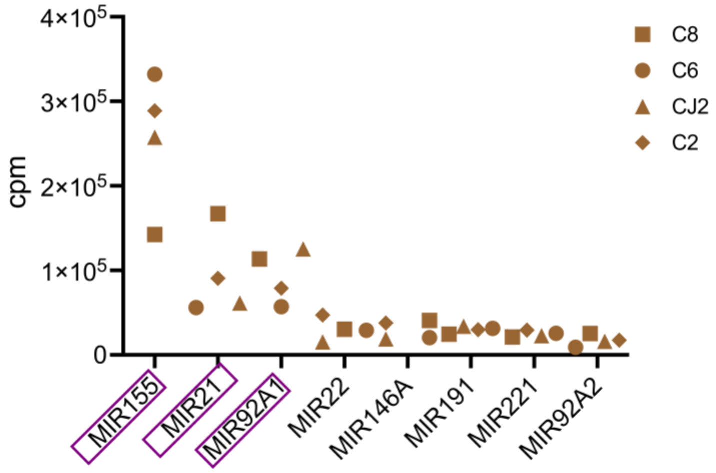


Supplementary Figure 9: Top miRNAs in sEV from HTLV/T with stimulatory effect on osteoclast differentiation. Purple boxes indicate most abundant species identified by small RNA sequencing.

### Supplementary Table 1: List of Primers Used

| For Genomic DNA qPCR | Forward 5'-3' | Reverse 5'-3' |
|---|---|---|
| Human *IL6* | ATTGGGAGCCCACACTCGAA | ATCACCTAGTCCACGCCCAA |
| Mouse *B2m* | ATGCTGAGGACCTTGTGAGC | GGAAGGCAGTAGGGAGAGGA |
| For RT-qPCR | | |
| Mus Musculas TRAP (*Acp5*) | CGACCATTGTTAGCCACATACG | TCGTCCTGAAGATACTGCAGGTT |
| Mus Musculas Cathepsin K (*Ctsk*) | ATATGTGGGCCAGGATGAAAGTT | TCGTTCCCCACAGGAATCTCT |
| Mus Musculas Calcitonin receptor (*CALCR*) | GCAACCGAACCTGGTCCAACTAT | AAGCAGCAATCGACAAGGAGTGA |
| Mus Musculas Dcstamp (*DCSTAMP*) | GACCTTGGGCACCAGTATTT | CAAAGCAACAGACTCCCAAA |
| Mus Musculas v-ATPase (*ATP6v0d2*) | AGTGCAGTGTGAGACCTTGG | TCTGCAGAGCTTCTTCCTCA |
| Human RANKL (*TNFSF11*) | GGCCACAGCGCTTCTCAG | GAGTGACTTTATGGGAACCCGAT |
| Human OPG (*TNFRSF11B*) | AGTCCGTGAAGCAGGAGTG | CCATCTGGACATTTTTTGCAAA |
| Human *GAPDH* | TGTGATGGGTGTGAACCACGAGAA | GAGCCCTTCCACAATGCCAAAGTT |
| Mus Musculas *Hprt1* | CCTAAGATGATCGCAAGTTG | CCACAGGGACTAGAACACCTGCTAA |
| Mus Musculas *GAPDH* | AGGTCGGTGTGAACGGATTTG | TGTAGACCATGTAGTTGAGGTCA |

### Supplementary Methods

#### Proteomics:

#### Sample Preparation

Exosomes isolated from human cell culture samples were solubilized in 35 µl of SDS buffer (4% (wt/vol)), 100 mM Tris-HCl pH 8.0) with sonication in a water bath sonicator (VWR, 150D) at RT for 10min at power level 9. Protein disulfide bonds were reduced using 100 mM dithiothreitol (DTT; Pierce, cat. no. 20291) with heating to 95 ºC for 10 min. Samples were digested as previously described (PMID 19377485). Reduced samples were mixed with 600 µl 100 mM Tris-HCL buffer, pH 8.5 containing 8 M urea (Sigma, cat no. U4884-500g) (UA buffer) and transferred to the top of a 30,000 molecular weight cut-off filter (Millipore, part# MRCF0R030) and spun in a microcentrifuge (Eppendorf) at 10,000 rcf for 10 min. An additional 300 µl of UA buffer was added and the filter was spun at 10,000 rcf for 10 min in a microcentrifuge. The flow through was discarded and the proteins were alkylated using 100 µl of 50 mM iodoacetamide (IAM, Pierce, Cat. No. A39271) in UA buffer. IAM in UA buffer was added to the top chamber of the filtration unit. The samples were gyrated at 550 rpm using a Thermomixer (Eppendorf) at room temperature for 30 min in the dark. The filter was spun at 10,000rcf for 10 min and the flow through discarded. Unreacted IAM was washed through the filter with two additions of 200 µl of UA buffer, and centrifugation at 10,000 rcf for 10 min after each buffer addition. The UA buffer was exchanged with digestion buffer (DB), 50 mM ammonium bicarbonate buffer, pH 8. Two sequential additions of DB (200 µl) with centrifugation after each addition to the top chamber was performed. The filters were transferred to a new collection tube and 100 µl DB containing 1

mAU of LysC (Wako Chemicals, cat. no. 129-02541) was added and samples were incubated at 37 °C for 2 h. Trypsin (1 µg) (Promega, Cat. No. V5113) was added and samples were incubated overnight at 37 °C. The filters were spun at 10,000 rcf for 15 min to collect the peptides in the flow through. The filter was washed with 50 µL 100mM ammonium bicarbonate buffer and the wash was collected with the peptides. Peptides were acidified with trifluoroacetic acid (TFA; Sigma, cat. no. 91707) to a final concentration of 1% (vol/vol) and were desalted using two micro-tips (porous graphite carbon, BIOMETNT3CAR) (Glygen) on a Beckman robot (Biomek NX) (Chen PMID 22338125). The peptides were eluted with 60% (vol/vol) acetonitrile (MeCN; J.T. Baker, cat. no. 9829-03) in 0.1% (vol/vol) TFA and dried in a Speed-Vac (Thermo Scientific, Model No. Savant DNA 120 concentrator). Samples were dissolved in 20 µl of 1% (vol/vol) MeCN in water. An aliquot (10%) was removed for quantification using the Pierce Quantitative Fluorometric Peptide Assay kit (Thermo Scientific, Cat. No. 23290).
The remaining peptides were transferred to autosampler vials (Sun-Sri, Cat. No. 200046), dried and stored at -80 °C.

UPLC-timsTOF-MS.
The peptides were analyzed using trapped ion mobility time-of-flight mass spectrometry (PMID3038 5480). Peptides were separated using a nano-ELUTE  chromatograph (Bruker Daltonics. Bremen, Germany) interfaced to a timsTOF Pro mass spectrometer (Bruker Daltonics) with a modified nano-electrospray source (CaptiveSpray, Bruker Daltonics). The mass spectrometer was operated in PASEF mode (PMID30385480). The samples in 2 µl of 1% (vol/vol) formic acid (FA; Sigma-Aldrich, cat. no. 56302) were injected onto a 75 µm i.d. × 25 cm Aurora Series column with a CSI emitter (Ionopticks). The column temperature was set to 50 °C. The column was equilibrated using constant pressure (800 bar) with 8 column volumes of solvent A (0.1% (vol/vol) FA. Sample loading was performed at constant pressure (800 bar) at a volume of 1 sample pick-up volume plus 2 µl. The peptides were eluted using one column separation mode with a flow rate of 300 nl/min and using solvents A (0.1% (vol/vol) FA) and B (0.1% (vol/vol) FA/MeCN): solvent A containing 2% B increased to 17% B over 60 min, to 25% B over 30 min, to 37% B over 10 min, to 80% B over 10 min and constant 80% B for 10 min. The MS1 and MS2 spectra were recorded from m/z 100 to 1700.
The collision energy was ramped stepwise as a function of increasing ion mobility: 52 eV for 0–19% of the ramp time; 47 eV from 19–38%; 42 eV from

38–57%; 37 eV from 57–76%; and 32 eV for the remainder. The TIMS elution voltage was calibrated linearly using the Agilent ESI-L Tuning Mix (m/z 622, 922, 1222).

MS data analysis. The MS2 spectra from peptides with +2, +3 and +4 charge states were analyzed using Mascot software (Perkins DN, Pappin DJ, Creasy DM, Cottrell JS, PMID: 10612281) (Matrix Science, London, UK; version 2.8.0.1). Mascot was set up to search against a UniProt reference databases of human proteins (20,512 entries, downloaded February 2021) and common contaminant proteins (cRAP, v1.0 Jan. 2012; 116 entries), assuming the digestion enzyme was trypsin with a maximum of 4 missed cleavages allowed. The searches were performed with a fragment ion mass tolerance of 20 ppm and a parent ion tolerance of 20 ppm. Carbamidomethylation of cysteine was specified in Mascot as a fixed modification. Deamidation of asparagine, deamidation of glutamine, pyro-glutamate formation from n-terminal glutamine, acetylation of protein N-terminus and oxidation of methionine were specified as variable modifications. Peptide spectrum matches (PSM) were filtered at 1% false-discovery rate (FDR) by searching against a reversed database and the ascribed peptide and protein identities were accepted.

The processing, quality assurance and analysis of LC-MS data were performed with proteoQ (version 1.5.0.0, https://github.com/qzhang503/proteoQ), software developed with the tidyverse approach ((i) Hadley Wickham. Advanced R, Second Edition (Chapman & Hall/CRC The R Series) (ii) Hadley Wickham (2017). tidyverse: Easily Install and Load the 'Tidyverse'. R package version 1.2.1. https://CRAN.R-project.org/package=tidyverse) with open source software for statistical computing and graphics, R (R Core Team (2021). R: A language and environment for statistical computing. R Foundation for Statistical Computing, Vienna, Austria. URL https://www.R-project.org/). and RStudio (RStudio Team (2016). RStudio: Integrated Development for R. RStudio, Inc., Boston, MA URL http://www.rstudio.com/). The precursor intensities were converted to logarithmic ratios (base 2), relative to the average precursor intensity across all samples. Within each sample, Dixon's outlier removals were carried out recursively for peptides with greater than two identifying PSM's. The median of the ratios of PSM that could be assigned to the same peptide was first taken to represent the ratios of the incumbent peptide. The median of the ratios of peptides was then taken to represent the ratios of the inferred protein. To align protein ratios across samples, likelihood functions were first estimated for the log-ratios of proteins using finite mixture modeling, assuming two-component Gaussian mixtures (

Tatiana Benaglia, Didier Chauveau, David R. Hunter, Derek Young (2009). mixtools: An R Package for Analyzing Finite Mixture Models. Journal of Statistical Software, 32(6), 1-29. URL http://www.jstatsoft.org/v32/i06/). The ratio distributions were then aligned so that the maximum likelihood of log-ratios was centered at zero for each sample. Scaling normalization was performed to standardize the log-ratios of proteins across all samples. To reduce the influence of outliers from either log-ratios or reporter-ion intensities, the values between the 5th and 95th percentile of log-ratios and 5th and 95th percentile of intensity were used in the calculations of standard deviations.

Bioinformatic and statistical analysis.

Metric multidimensional scaling (MDS) and Principal component analysis (PCA) of protein log2-ratios was performed with the base R function stats:cmdscale and stats:prcomp, respectively. Heat-map visualization of protein log2-ratios was performed with heatmap ( Raivo Kolde (2019). pheatmap: Pretty Heatmaps. R package version 1.0.12. https://CRAN.R-project.org/package=pheatmap). Linear modelings were performed using the contrast fit approach in limma ( Ritchie, M.E., Phipson, B., Wu, D., Hu, Y., Law, C.W., Shi, W., and Smyth, G.K. (2015). limma powers differential expression analyses for RNA-sequencing and microarray studies. Nucleic Acids Research 43(7), e47.), to assess the statistical significance in protein abundance differences between indicated groups of contrasts. Adjustments of p-values for multiple comparison were performed with Benjamini-Hochberg (BH) correction.
